# Study of Metyltetraprole, an unusual agrofungicide targeting the Q_o_-site of cytochrome *bc*_1_ complex

**DOI:** 10.1101/2021.10.01.462775

**Authors:** Claudia Serot, Thomas Michel, Valentine Lucas, Brigitte Meunier

## Abstract

The mitochondrial respiratory chain *bc*_1_ complex is a proven target of agrofungicides. Most of them are Q_o_-site antagonists (*i*.*e* QoIs), competing with the substrate ubiquinol, and likely share the same binding mode as the widespread Q_o_-site resistance mutation G143A confers cross-resistance. Metyltetraprole (MTP) presents an exception as studies with phytopathogenic fungi showed that the inhibitor was unaffected by G143A. Here, we used the yeast model to investigate its mode of action. Analysis of *bc*_1_ complex mutants supports a Q_o_-site binding for MTP. However the compound seems distinct to other QoIs, such as azoxystrobin, in various ways, namely; 1) G143A was without effect on MTP, as previously reported. 2) The level of MTP resistance of mutants was higher in *bc*_1_ complex activity assays than in growth assays while the opposite was observed with azoxystrobin. 3) Steady-state kinetics used to characterise the mode of action of MTP also revealed differences compared to other QoIs.

## Introduction

Many fungal plant pathogens rely on respiration for cell maintenance and proliferation and thus require an active respiratory chain. The mitochondria cytochrome *bc*_1_ complex is a key component of the respiratory chain. Inhibitors targeting that enzyme could be efficient tools for pathogen control.

The *bc*_1_ complex is a homodimeric complex anchored in the inner mitochondrial membrane that functions as a protonmotive ubiquinol:cytochrome *c* oxidoreductase, conserving and converting the energy obtained from the oxidation of ubiquinol by cytochrome *c* into a transmembrane proton gradient which can be used for other processes, such as ATP synthesis by the F_o_F_1_ ATP synthase. The monomer of the eukaryote enzyme consists of 10-11 discrete subunits, encoded by nuclear genes, with the exception of one subunit, cytochrome *b*, which is encoded by a mitochondrial gene. The highly conserved catalytic core of the enzyme is formed by three subunits bearing the redox prosthetic groups, cytochrome *b*, cytochrome *c*_1_ and the Rieske iron-sulphur protein. The membrane-spanning cytochrome *b* provides the two ubiquinol/ubiquinone binding sites of the enzyme, located on opposite sides of the membrane, and termed Q_o_ and Q_i_ sites. These two sites are also the binding sites of inhibitors.

Most of the agrofungicides on the market that target the *bc*_1_ complex are Q_o_-site inhibitors (QoIs). They include synthetic strobilurins such as azoxystrobin. These agrofungicides likely share the same binding mode as the same resistance mutation in the Q_o_-site, G143A, confers cross-resistance to all of these inhibitors. Unfortunately, G143A causing high levels of resistance is widespread in pathogen fungi, compromising the efficiency of the whole family of QoIs in disease control. Inspection of the atomic structure of the *bc*_1_ complex with bound azoxystrobin, for example, shows that replacement of glycine by alanine causes steric clashes with the inhibitor MOA-bearing phenyl moiety.

Metyltetraprole (MTP) is a novel QoI that does not seem affected by the resistance mutation G143A [1-3]. The compound has a side-chain similar to that of the synthetic strobilurin pyraclostrobin, but has a unique tetrazolinone-moiety. It was suggested that the tetrazolinone-moiety would not form the same highly specific interactions with the Q_o_-site as do other strobilurin-based QoIs but could accommodate changes in the target. The steric hindrance caused by G143A that compromises the binding of QoIs would be limited by the unique size and shape of the tetrazolinone. The atomic structure of WT and G143A *bc*_1_ complex with bound MTP is needed to confirm its distinct mode of binding with its target. Meanwhile, the study of other Q_o_-site mutations on MTP sensitivity could provide information on its binding mode. In this report, we used yeast *Saccharomyces cerevisiae* as model to investigate MTP mode of action. High resolution atomic structures of the *bc*_1_ complex co-crystallised with a variety of Q_o_-site occupants are available for the yeast enzyme and also for chicken, bovine and bacterial *bc*_1_ complexes (for instance see survey in [4]), but not yet for phytopathogen fungi enzymes. Yeast and phytopathogenic fungi cytochrome *b* present a high degree of sequence identity, making yeast a relevant model. In addition, yeast being amenable to mitochondrial transformation, engineered mutations can be introduced into the mitochondrially-encoded cytochrome *b* gene. Activity of the *bc*_1_ complex can be easily measured in crude mitochondrial membrane preparations from WT and mutants.

Here we studied the effect of Q_o_-site mutations in yeast *bc*_1_ complex on MTP inhibition. Our data clearly supports a Q_o_-site binding for MTP. However, the compound seems to have an unusual mode of inhibition, different to other QoIs such as azoxystrobin.

## Materials and Methods

### Materials and growth media

Cytochrome *c*, decylubiquinone and azoxystrobin were obtained from Sigma Aldrich. Metyltetrapole was provided by Sumitomo Chemical. The following media were used for yeast growth: YPD (1% yeast extract, 2% peptone, 3% glucose), YPGal (1% yeast extract, 2% peptone, 2% glycerol, 0.1% glucose), YPG (1% yeast extract, 2% peptone, 2% glycerol) and YPEth (1% yeast extract, 2% peptone, 2% ethanol).

### Yeast strains

The mutations were introduced in yeast cytochrome *b* gene as described in [5,6]. In all experiments, control and mutants have identical nuclear and mitochondrial genomes with the exception of the mutations in cytochrome *b* gene.

### Growth assays

Yeast cells were grown in 5 mL YPEth with increasing concentration of inhibitors. Cultures were inoculated at an OD_600nm_ of 0.2 and incubated at 28°C with vigorous shaking for three days. OD_600 nm_ were then measured. The experiments were repeated at least twice and the data averaged. The IC_50s_ (mid-point inhibition concentrations) were estimated from the plots of final OD_600nm_ versus inhibitor concentation.

### NADH- and decylubiquinol-cytochrome *c* reductase activity

Yeast mitochondria were prepared as in [7]. Briefly, yeast grown in YPGal medium were harvested at mid-log phase. Protoplasts were obtained by enzymatic digestion of the cell wall using zymolyase in an osmotic protection buffer. Mitochondria were then prepared by differential centrifugation following osmotic shock of the protoplasts. Mitochondrial samples were aliquoted and stored at −80°C. Concentration of *bc*_1_ complex in the mitochondrial samples was determined from dithionite-reduced optical spectra, using ε=28.5 mM^-1^ cm^-1^ at 562 nm *minus* 575 nm. Cytochrome *c* reductase activities were determined at room temperature by measuring the reduction of cytochrome *c* (final concentration of 20 µM) at 550 nm *minus* 540 nm over 1-min time-course in 10 mM potassium phosphate pH 7 and 1 mM KCN (to inhibit cytochrome *c* oxidase activity). Lauryl-maltoside (0.01% w/v) was added to the reaction buffer for the decylubiquinol-cytochrome *c* reduction assays. Mitochondria were added to obtain a final concentration of around 6 nM *bc*_1_ complex. The reaction was initiated by the addition of 200 μM NADH or 40 μM decylubiquinol and initial rates were measured. Sensitivity to inhibitors was tested by adding increasing concentrations of inhibitors to the reaction mixture. The measurements were repeated at least twice and averaged. IC_50_ values (mid-point inhibition concentrations) were estimated from the inhibition curves.

### Preparation of mitochondrial samples from yeast and *Zymoseptoria tritici*

The *Z. tritici* WT (IPO323) and G143A mutant were obtained from A-S Walker (INRAE, Grignon, France). The cells were grown for 5 days at 17°C in 250 mL YPD medium with vigorous agitation. The cultures were then harvested and washed. The pellets were frozen in liquid nitrogen and ground. The mitochondrial were then prepared from the ground cells by following the method used for yeast mitochondrial samples [7].

## Results & Discussion

### 1. Effect of the QoI resistance mutation G143A on MTP sensitivity

The most widespread target site mutation G143A causes a high level of QoI resistance in field pathogen fungi. We introduced this mutation into yeast cytochrome *b* and tested its impact on MTP sensitivity in respiratory growth (YPEth medium) and *bc*_1_ complex activity assays. By comparison, the effect of G143A on azoxystrobin sensitivity was also examined (Table 1).

**Table 1.** Effect of G143A on MTP and azoxystrobin sensitivity of yeast growth and *bc*_1_ complex activity

| inhibitor/strain | growth |  | *bc*<sub>1</sub> complex activity |  |
| --- | --- | --- | --- | --- |
|  | IC<sub>50</sub> (nM) | RF | IC<sub>50</sub> (nM) | RF |
| MTP |  |  |  |  |
| WT | 47 | 1 | 1.5 | 1 |
| G143A | 30 | 0.6 | 1 | 0.7 |
| Azoxystrobin |  |  |  |  |
| WT | 60 | 1 | 35 | 1 |
| G143A | > 15 000 | > 250 | > 10 000 | > 280 |

Growth and *bc*<sub>1</sub> complex assays were performed as described in Materials and Methods. The IC<sub>50</sub>s, mid-point inhibition concentrations, were estimated from the inhibitor titration curves. Each measurement was repeated at least twice and the values averages. Standard errors were ≤ 15% of the measured values. RF, resistance factor (IC<sub>50</sub> mutant/IC<sub>50</sub> WT).

G143A, as expected, dramatically decreased azoxystrobin sensitivity whereas the mutation had no effect on MTP sensitivity. This is in agreement with previous report showing that G143A had only a mild effect on the sensitivity of *Zymoseptoria tritici* (and other plant pathogen fungi) in enzymatic assays [2]. To further confirm that observation, we prepared mitochondria from *Z*.*tritici* control and G143A mutant strains and tested the QoI sensitivity of the *bc*_1_ complex (Table 2). The RFs obtained here were very similar to the values previously reported [2]. Clearly, both in *Z*.*tritici* and yeast, MTP efficiency was not or very mildly affected by G143A.

**Table 2.**
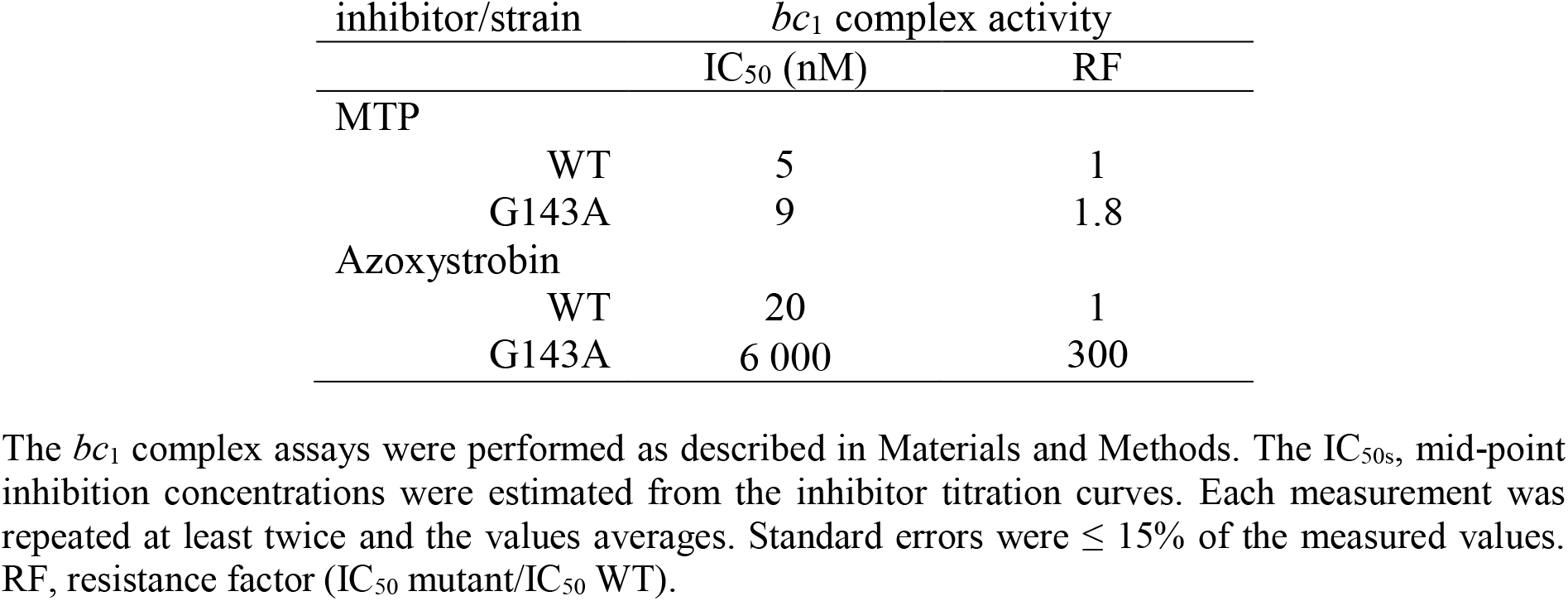
Effect of G143A on MTP and azoxystrobin sensitivity of *Z*.*tritici bc*_1_ complex activity

| inhibitor/strain | *bc*<sub>1</sub> complex activity |  |
| --- | --- | --- |
|  | IC<sub>50</sub> (nM) | RF |
| MTP |  |  |
| WT | 5 | 1 |
| G143A | 9 | 1.8 |
| Azoxystrobin |  |  |
| WT | 20 | 1 |
| G143A | 6 000 | 300 |

The *bc*<sub>1</sub> complex assays were performed as described in Materials and Methods. The IC<sub>50</sub>s, mid-point inhibition concentrations were estimated from the inhibitor titration curves. Each measurement was repeated at least twice and the values averages. Standard errors were ≤ 15% of the measured values. RF, resistance factor (IC<sub>50</sub> mutant/IC<sub>50</sub> WT).

### 2. Effect of Q_o_-site mutations on MTP sensitivity

We then tested, in yeast, the effect of substitutions of four other residues involved in QoI sensitivity/resistance, namely F129, Y132, G137 and L275, and of another substitution of residue 143.

The substitution F129L was found in field isolates of several fungi species (see https://www.frac.info/home). The mutations F129S, Y132C and G143S were observed in *Peronophythora litchii* after treatment with QoIs in the laboratory [8]. G137R was observed in isolates of *Pyrenophora tritici-repentis* [9]. L275S and L275F were first identified in yeast after selection on the myxathiozol [10]. We generated yeast mutants harbouring F129L/S, Y132C, G143S or L275S/F and two additional mutants, F129G (to test a smaller residue) and Y132F (a conservative substitution as a control).

First we performed a growth assay on plates to test the respiratory growth competence and sensitivity of the mutants to MTP and azoxystrobin (Fig.1). All the mutants grew well on glucose medium (glucose can be used for energy supply by both fermentation and respiration). Y132C and G143S showed a weaker growth on glycerol (glycerol, as ethanol, is exclusively used for energy production by respiration). Growth test on inhibitor supplemented media showed different pattern of resistance. F129G presented a marked resistance to MTP; F129L and F129S showed a moderate resistant to MTP. The level of resistance to azoxystrobin was higher in F129L and F129G than in F129S. Y132C grew on MTP-but not on azoxystrobin-supplemented media. The change Y132F had no effect. G143A, highly resistant to azoxystrobin, remained sensitive to MTP, whereas Y143S was resistant to both inhibitors. Finally, L275F showed a weak growth on MTP medium and none on azoxystrobin, while L275S demonstrated resistance to both QoIs. G137R was excluded from this test as the mutant grow very poorly on respiratory medium.

**Figure 1.**
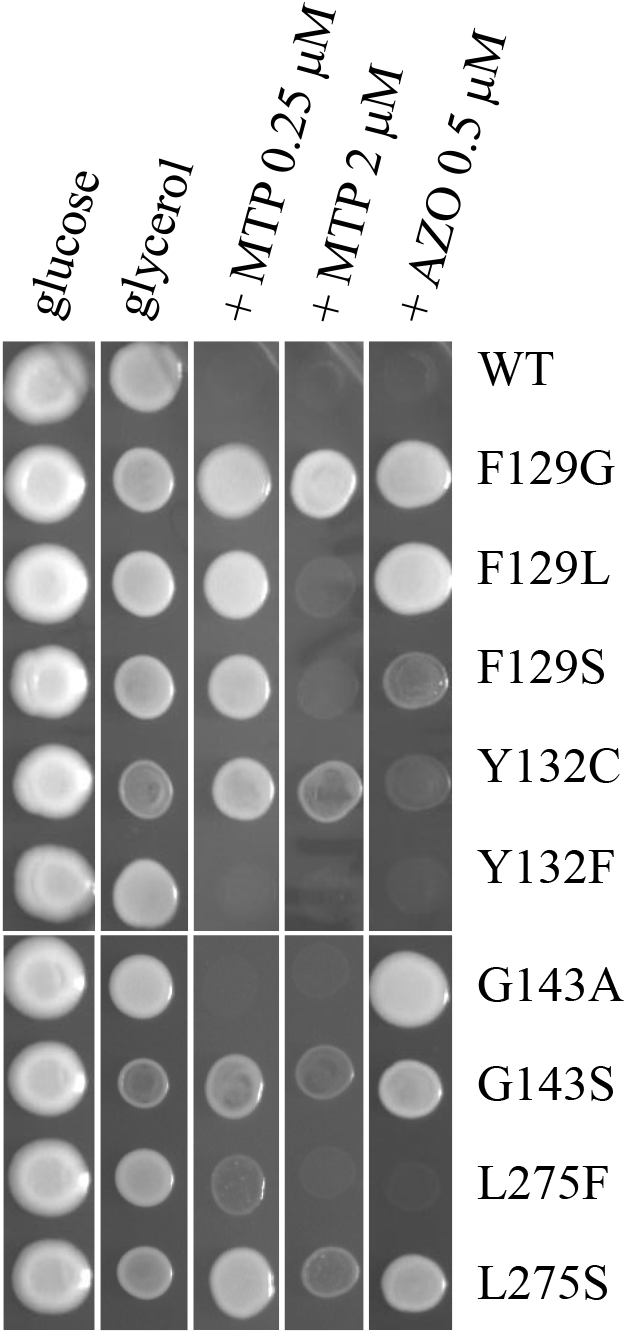
Effect of Q_o_-site mutations on growth sensitivity to MTP and azoxystrobin. Suspensions in water of cells pre-grown on glucose plates were spotted on plates containing either glucose (fermentative medium, YPD) or glycerol (respiratory medium, YPG) with/without MTP or azoxystrobin (AZO) and incubated for three days at 28 °C.

The study was pursued with the MTP-resistant mutants. We checked the respiratory growth competence of the mutants in YPEth and determined IC_50s_ (concentrations of MTP required to obtain 50% inhibition of cell growth) and resistance factors (RF, as IC_50_ mutant / IC_50_ WT). In parallel, we prepared mitochondria and tested the *bc*_1_ complex activity and sensitivity to MTP (Table 3).

**Table 3.** Growth and *bc*_1_ complex sensitivity to MTP of WT and Q_o_-site mutants

| strain | growth |  |  | *bc*<sub>1</sub> complex activity |  |  |
| --- | --- | --- | --- | --- | --- | --- |
|  | final OD<sub>600nm</sub><br>(% WT) | IC<sub>50</sub><br>(nM) | RF | activity (s<sup>-1</sup>)<sup>a</sup><br>(% WT) | IC<sub>50</sub> /<br>[*bc*<sub>1</sub>]<sup>b</sup> | RF |
| WT | 100 | 47 | 1 | 100 | 0.3 | 1 |
| F129G | 78 | 900 | 19 | 105 | 40 | 133 |
| F129L | 83 | 275 | 6 | 95 | 15 | 51 |
| F129S | 83 | 250 | 5 | 100 | 10 | 32 |
| Y132C | 26 | 500 | 11 | 14 | 21 | 69 |
| G137R | nd | nd | nd | 26 | 0.2 | 0.7 |
| G143S | 33 | 250 | 5 | 20 | 4 | 13 |
| L275S | 84 | 250 | 5 | 45 | 31 | 104 |

Growth and *bc*<sub>1</sub> complex assays were performed as described in Materials and Methods. Each measurement was repeated at least twice and the values averages. Standard errors were ≤ 20% of the measured values for the growth assays and ≤ 10% for the *bc*<sub>1</sub> complex assays. IC<sub>50</sub>, mid-point inhibition concentration. RF, resistance factor (IC<sub>50</sub> mutant/IC<sub>50</sub> WT). <sup>a</sup> The *bc*<sub>1</sub> complex activities are measured as cytochrome *c* reduced per *bc*<sub>1</sub> complex and per second using 40 μM decylubiquinol as electron donor. The WT activity was 65 s<sup>-1</sup>. <sup>b</sup> The IC<sub>50</sub> values are presented as IC<sub>50</sub> reported on the concentration of *bc*<sub>1</sub> complex.

The changes Y132C and G143S resulted in a growth defect (already observed in Fig 1) and a marked decrease in *bc*_1_ complex activity. L275S caused a two-fold decrease in *bc*_1_ complex activity but with no effect on growth competence. G137R impaired *bc*_1_ complex activity leading to a very severe defect in respiratory growth as previously reported [11]. The substitutions of residue 129, F129G/L/S, had no effect on growth competence and enzymatic activity.

Both growth and activity assays confirmed the resistance to MTP conferred by mutations F129G/L/S, Y132C, G143S and L275S. G137R did not cause resistance. As a control, we monitored the impact of two Q_i_-site mutations, namely G37V and L198F that confer resistance to QiIs. These Q_i_-site mutations had no effect on MTP sensitivity (not shown).

To broaden the mutational analysis of the Q_o_-site, we also checked the effect of substitutions of residues E272 and Y279 in a growth test as in Fig.1. It was shown that QoIs may form hydrogen-bonds with the backbone amide moiety of E272 [12]. Mutations E272V/T and S had no effect on MTP sensitivity (not shown). Y279C/S were found as atovaquone resistance mutations in *Plasmodium falciparum* and caused high level of atovaquone resistance in yeast [13]. These two mutations conferred also MTP resistance (not shown).

Thus all together, these data strongly support a Q_o_-site binding for MTP, as expected from its structural similarity with other QoIs, especially pyraclostrobin and from assays of binding competition between MTP and pyraclostrobin that suggested that the two compounds share the same binding site [1]. The resolution of the atomic structure of MTP-bound *bc*_1_ complex would be most informative. It could be expected that comparison with the structures of pyraclostrobin and other QoIs-bound enzymes would reveal similarities but also clear differences as G143A did not confer cross-resistance to MTP.

### 3. Unusual behaviour of MTP

As shown in Table 3, the growth assays revealed a moderate resistance to MTP, with RFs of 19 for F129G, 11 for Y132C and 5-6 for the remaining mutants. When the *bc*_1_ complex activities were tested, a marked resistance was observed, with RF up to 133 and 104 for F129G and L275S, respectively.

To confirm the differences in RFs between enzymatic activity assays and growth assays, we monitored the NADH-cytochrome *c* reductase activity of WT and mutants F129G, F129L, F129S and L275S. The mutants Y132C, G137R and G143S presenting a marked decrease in activity were not studied further. Table 4 compared the IC_50s_ and RFs obtained by monitoring decylubiquinol-cytochrome *c* reduction activity (*bc*_1_ complex activity, data from Table 3) with those obtained by monitoring the NADH-cytochrome *c* reduction activity. In these latter assays, we measured the combined activity of NADH-dehydrogenase (NADH-ubiquinone oxidoreductase) and *bc*_1_ complex. Thus no exogenous quinol was added. Note that MTP had no effect on NADH-dehydrogenase activity itself (measured up to 10 μM). Interestingly, the IC_50s_ of the mutants were two to three-fold lower when using NADH as electron donor instead of decylubiquinol whereas the IC_50_ of the WT was unchanged, which resulted in decreased RFs. As a control, we also measured the succinate-cytochrome *c* reduction activities of WT and F129L (5mM succinate being used as electron donor instead of NADH). The same RF of 12 was obtained (data not shown).

**Table 4.** Sensitivity to MTP of WT and Q_o_-site mutants: comparison between decylubiquinol - and NADH-cytochrome *c* reductase activity

| strain | decylubiquinol-cytc reductase<sup>a</sup> |  | NADH-cytc reductase |  |
| --- | --- | --- | --- | --- |
|  | IC<sub>50</sub> / [*bc*<sub>1</sub>] | RF | IC<sub>50</sub> / [*bc*<sub>1</sub>] | RF |
| WT | 0.3 | 1 | 0.4 | 1 |
| F129G | 40 | 133 | 20 | 49 |
| F129L | 15 | 51 | 4.8 | 12 |
| F129S | 10 | 32 | 4.5 | 11 |
| L275S | 31 | 104 | 9.5 | 23 |

The impact of the Q_o_-site mutations on MTP binding might be weaker with the endogenous quinol than with the synthetic quinol analog, decylubiquinol, given the differences in the aliphatic/alkenyl sidechains of these two substrates.

As a comparison, we determined the IC_50s_ of azoxystrobin and RFs for three mutants, namely F129G, F129L and L275S that showed a clear resistance to that inhibitor with little or no growth defect (Fig.1, Table 3). The mutants presented a moderate resistance in *bc*_1_ complex assays (RFs of 3-8) and a two-fold - or more-higher resistance in growth assays (Table 5). This increased resistance of growth as compared to *bc*_1_ complex activity has been previously observed for other inhibitors, such as HDQ [14] and atovaquone [11]. Thus, MTP seems to have an unusual behaviour in that regard.

**Table 5.** Growth and *bc*_1_ complex sensitivity to azoxystrobin of WT, F129G, F129L and L275S

| strain | growth |  | *bc*<sub>1</sub> complex activity |  |
| --- | --- | --- | --- | --- |
|  | IC<sub>50</sub> (nM) | RF | IC<sub>50</sub> / [*bc*<sub>1</sub>] | RF |
| WT | 60 | 1 | 8 | 1 |
| F129G | >1,000 | >15 | 60 | 8 |
| F129L | >1,000 | >15 | 62 | 8 |
| L275S | 400 | 6.5 | 23 | 3 |

Growth and *bc*<sub>1</sub> complex assays were performed as described in Materials and Methods. Each measurement was repeated at least twice and the values averages. Standard errors were $\leq 20\%$ of the measured values for the growth assays and $\leq 10\%$ for the *bc*<sub>1</sub> complex assays. IC<sub>50</sub>, mid-point inhibition concentration. RF, resistance factor (IC<sub>50</sub> mutant/IC<sub>50</sub> WT).

Figure 2 summarises the differences in RFs and highlights the difference between MTP and azoxystrobin. It shows the RFs estimated from the activity assays (azoxystrobin resistance of decylubiquinol-cytochrome *c* reductase activities and MTP resistance of decylubiquinol- and NADH-cytochrome *c* reductase activities) reported to the RFs estimated from the growth assays.

**Figure 2.**
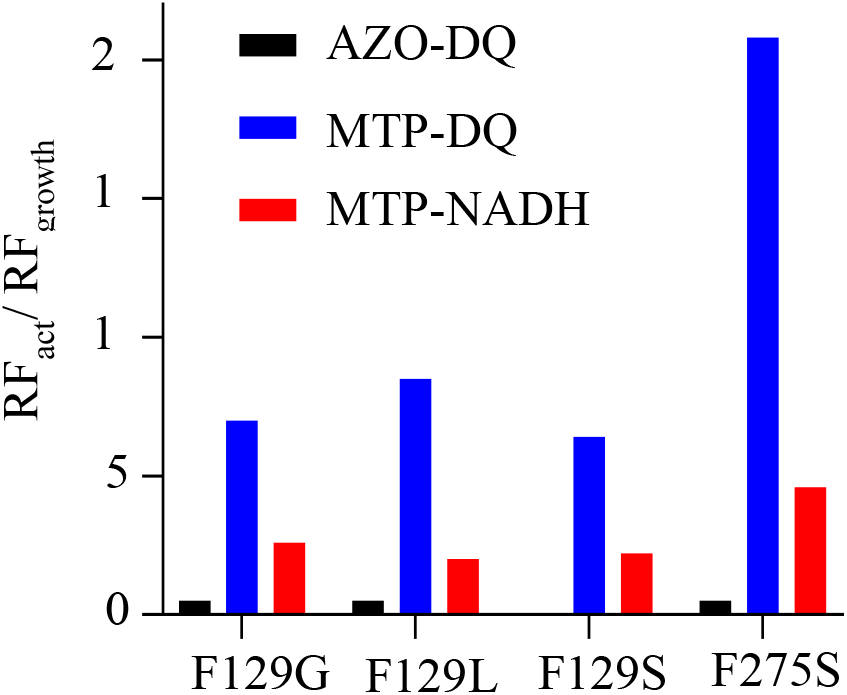
Comparison of the resistance factors. Resistance factors (RF) obtained from activity assays (RF_act_) were reported to RFs obtained from growth assays (RF_growth_). Decylubiquinol-cytochrome *c* reductase assays, resistance to azoxystrobin (AZO-DQ). Decylubiquinol- and NADH-cytochrome *c* reductase assays, resistance to MTP (MTP-DQ and MTP-NADH). Data are from Tables 3, 4 and 5.

To pursue the comparison between MTP and other QoIs, we performed assays to assess the mode of inhibition of MTP in WT and F129G. That mutant was chosen as it presented a high resistance to MTP with no decrease in *bc*_1_ complex activity.

Steady-state kinetics of the *bc*_1_ complex were previously used to characterise the inhibition mode of other compounds. Atovaquone and stigmatellin were showed to act as a competitive inhibitor of decylubiquinol [15]. The compounds bind in a region of the Q_o_-site in close proximity to the mobile domain of iron-sulphur protein, forming a hydrogen bond to a histidine ligand of the iron-sulphur protein [2Fe-2S] cluster, and restricting its movement [4,12]. These inhibitors are classed as *b*_L_ distal inhibitors (QoD-I), i.e. binding at a distal position within the Q_o_-site with respect to haem *b*_L_.

Methoxyacrylate (MOA) inhibitors oudmansin A, strobilurin A and MOA-stilbene were reported to act as non-competitive inhibitors [16]. But re-examining of the data indicated a mixed type inhibition, which was confirmed for MOA-stilbene [17]. A mixed type of inhibition would support a model of overlapping binding sites for MOA-stilben and ubiquinol (decylubiquinol was used in that study) [17], which would be in agreement with the structural analysis. The MOA inhibitors are classed as *b*_L_-proximal inhibitors (QoP-I). This class contains the synthetic strobilurins, such as azoxystrobin and pyraclostrobin. Because of its structural similarity with pyraclostrobin, MTP is expected to be a QoP-I.

To check whether MTP would show the same mode of inhibition, we monitored the *bc*_1_ complex activity using increasing concentrations of decylubiquinol and at different concentrations of MTP and observed the effect of MTP on V_max_ and K_m_ (Fig.3). Interestingly MTP appears to act as an uncompetitive inhibitor since both V_max_ and K_m_ decreased (Fig. 3A and B). An uncompetitive mode of inhibition would mean in theory that MTP binds to the enzyme-substrate complex (and not to the free enzyme), thus when decylubiquinol is bound at the Q_o_-site, which (for steric reasons) seems unlikely based upon the structure. The same test was then realised with the F129G mutant and a mixed inhibition was observed, as V_max_ decreased while K_m_ increased (Fig.3C), resembling the mode of inhibition observed for MOA-stilbene [17]. Thus apparently a different mode of inhibition was observed for the WT and mutant enzymes in that assay. This could perhaps be a clue to understand the difference in RFs (Fig.2).

**Figure 3.**
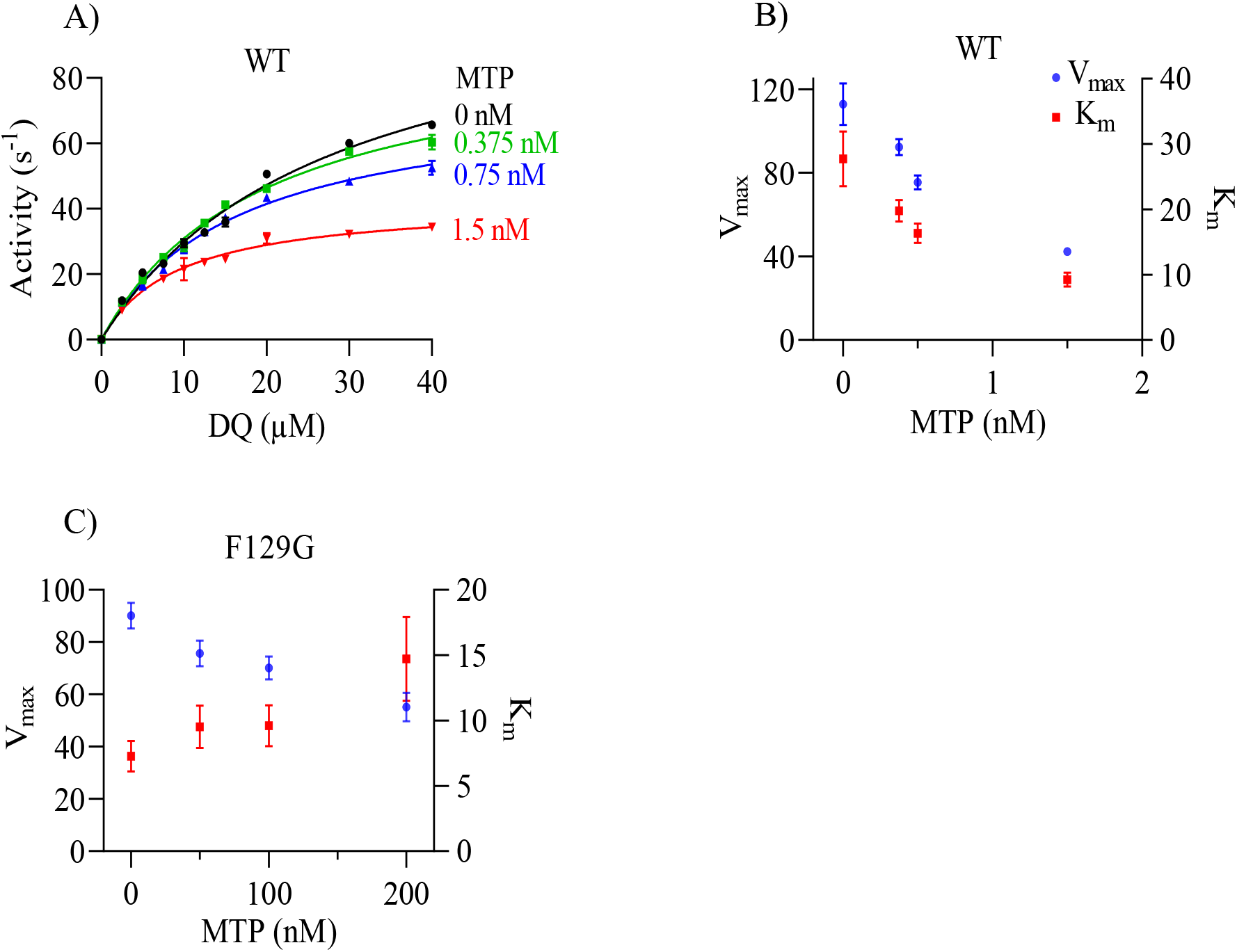
Effect of MTP on *bc*_1_ complex steady-state kinetics. A) Plot of WT *bc*_1_ complex activity *versus* decylubiquinol concentrations at different concentrations of MTP. The activities were monitored as described in Materials and Methods, using 2.5 to 40 μM decylubiquinol. Each measurement was repeat at least twice and the data averaged. The error bars represent the standard deviations. Curve fitting was performed with GraphPad Prism. B) Effect of MTP concentrations on V_max_ and K_m_ of the WT enzyme. Estimation of V_max_ and K_m_ was performed with GraphPad Prism using the data shown in A. The error bars represent the fitting errors. C) Effect of MTP concentrations on V_max_ and K_m_ of F129G enzyme. Data were obtained as in A.

Inhibitor titration of steady-state kinetics might not be appropriate when performed with a membrane bound enzyme, such as the *bc*_1_ complex and highly hydrophobic substrate and inhibitors, and tight binding inhibitors. Therefore one needs to be cautious when interpreting the data. Nevertheless, the results suggest that MTP present a peculiar behaviour, distinct from MOA-stilbene (and possibly other QoIs) that would warrant more detailed analysis. It might open new questions on single *versus* double occupancy of the Q_o_-site. The determination of the atomic structures of MTP-bound enzymes, WT and mutants, would probably reveal an unusual mode of binding.

## Conclusions

MTP appears as an unusual QoI. As previously observed with phytopathogenic fungi, G143A that compromises the efficiency of other QoIs was without effect on MTP, both in growth and *bc*_1_ complex activity assays.

The second most common QoI resistance mutation, namely F129L, had only a moderate effect on MTP inhibition as we observed a RF of 6 in growth assays and 12 in NADH- and succinate-cytochrome *c* reduction activities assays. By comparison, the RFs reported for the plant pathogen fungus *Pyrenophora teres* bearing the mutation F129L was 1.6 in growth assays [3] and 5.4 in succinate-cytochrome *c* reductase activities assays [2]. The (small) 2-4 fold difference in RFs between yeast and *P*.*teres* could be due to small differences in the Q_o_-site local structure that affect the impact of the mutation on MTP binding. It could be expected that a same mutation could have a different impact depending of the species. For example, growth sensitivity assays performed with various plant pathogen fungi bearing F129L revealed RFs for MTP between 0.7 and 3.3, and RFs for azoxystrobin between 18.5 and 204.3 [3]. We have previously observed that modification of the Q_o_-site by replacing a few residues of the yeast enzyme by their equivalents in humans or in *P*.*falciparum* changed its sensitivity towards inhibitors [18].

It would be useful to be able to predict which mutations could be a threat to the inhibitor potency. The chosen resistance mutations studied here, if they appeared in the fields, seem unlikely to jeopardise MTP efficiency, based on yeast data. F129L, F129S, G143S and L275S conferred a low level of resistance (RF of 5-6) in growth assays. Y132C resulted in a RF of 11 but was associated with growth defect. F129G caused the highest resistance level (RF of 19) but the amino-acid substitution requires two nucleotide changes at the codon for F129, which is unlikely to occur in the fields. The study of a broader series of yeast mutants might be useful. However the direct selection of resistant mutants of plant pathogen fungi would be more informative on the possible risk of resistance development.

## Acknowledgements

We are grateful to N. Fisher, Michigan State University, USA, for his helpful discussion of the results and corrections of the manuscript.

